# Integrative mapping of spatial transcriptomic and amyloid pathology in Alzheimer’s disease at single-cell resolution

**DOI:** 10.1101/2023.05.07.539389

**Authors:** Guang-Wei Zhang, Shangzhou Xia, Nicole K. Zhang, Fan Gao, Berislav V. Zlokovic, Li I. Zhang, Zhen Zhao, Huizhong W. Tao

**Author notes:** Correspondence should be addressed to Z. Zhao, or H.W.Tao. These authors contributed equally to this work (Guang-Wei Zhang, Shangzhou Xia).

## Abstract

Alzheimer’s disease (AD) is a complex neurodegenerative disorder that affects millions of people worldwide. Despite decades of research, the underlying molecular and cellular changes of AD remain unresolved, especially in terms of the spatial structure of gene expression changes that correlates with pathology, e.g. amyloid beta (A-beta) plaques. Recent advances in imaging-or sequencing-based single-cell spatial transcriptomics have allowed a systematic dissection of molecular and cell architectures in the brain and other tissues. In this study, we employed the recently developed Stereo-seq technology to spatially profile the whole-genome transcriptomics in the 5xFAD mouse model and established the methodology to analyze the specific neuronal transcriptomic changes spatially correlated with amyloid pathology at single cell resolution. More specifically, we developed a pipeline for integrative image- and non-image-based cell segmentation, VoxelMorph-based non-linear alignment, and Unet-based object detection to achieve reliable transcriptomics analysis at the single-cell resolution, and investigated the spatial relationship between diverse neuronal clusters and A-beta depositions. This work has demonstrated the potential of using the Stereo-seq technology as a powerful tool to investigate AD and other complex neurological disorders.

## Introduction

Alzheimer’s disease (AD) is the most prevalent neurodegenerative disorder that still lacks an effective treatment (Scheltens et al. 2016). Biomarkers, such as amyloid beta (Murphy and LeVine III 2010) and tau proteins (Iqbal et al. 2010), have been identified and are often used to track disease progression and evaluate potential therapies. Amyloid beta (A-beta) forms the hallmark amyloid plaques, while tau proteins become hyperphosphorylated and form neurofibrillary tangles in the brains of Alzheimer’s patients. Previous studies have suggested that these pathological changes exhibit specific spatial patterns in both human AD patients (Heiko Braak, Eva Braak, and Peter Kalus 1989) and animal models of AD (Gail Canter et al. 2019; Aldis P. Weible and Michael Wehr 2022). For example, neuropathology studies demonstrate that A-beta pathology exhibits a specific spatiotemporal pattern, starting in association cortices and spreading from neocortex to allocortex, which potentially drives tau pathology at later stages (van der Kant, Goldstein, and Ossenkoppele 2020). These observations suggest that revealing the spatial architecture of AD pathology may potentially provide additional clues to the selective vulnerability in AD.

In recent years, single-cell RNA sequencing has emerged as a powerful technique for investigating gene expression changes at the single-cell level. This technology allows researchers to measure gene expression in individual cells, providing unprecedented insights into cell diversity and functions. Previous studies have adopted the single-cell RNA sequencing (scRNA-seq) technique to characterize the transcriptomic changes in patients diagnosed with AD (Mathys et al. 2019; Liu-Lin Xiong et al. 2021; Trygve E. Bakken et al. 2018; Marta Olah et al. 2020; Blue B. Lake et al. 2016; Jorge L. Del-Aguila et al. 2019), as well as in animal models such as the 5xFAD (Yaming Wang et al. 2016; Habib et al. 2020; Minghui Wang et al. 2022) and APP/PS1 mice (Minghui Wang et al. 2022; Emma Gerrits et al. 2021; Soyon Hong et al. 2016). These studies generated fruitful results and offered a considerably deeper understanding of the cellular and molecular changes associated with AD. However, the scRNA-seq data cannot provide information on the location of individual cells within the tissue or the spatial relationship of individual cells with pathological hallmarks, such as plaques and tangles in AD.

Spatial transcriptomics is a rapidly developing field (Moses and Pachter 2022; Rao et al. 2021), and has been quickly adapted in neuroscience (Cameron G Williams et al. 2022; Jennie L. Close, Brian R. Long, and Hongkui Zeng 2021; Ed S. Lein, Lars E. Borm, and Sten Linnarsson 2017; Jürgen Germann et al. 2022; Patrik L. Ståhl et al. 2016), developmental (Cameron G Williams et al. 2022; Ed S. Lein, Lars E. Borm, and Sten Linnarsson 2017; Jürgen Germann et al. 2022; Patrik L. Ståhl et al. 2016) and cancer biology (Taku Monjo et al. 2022; Rao et al. 2021; Niyaz Yoosuf et al. 2020; Emelie Berglund et al. 2018; Alona Levy-Jurgenson et al. 2020; Kim Thrane et al. 2018). A recent study has investigated molecular changes and cellular interactions around amyloid plaques using spatial transcriptomics in an AD mouse model. It has revealed early alterations in a gene co-expression network enriched for myelin and oligodendrocyte genes, and late-stage multicellular co-expression networks of plaque-induced genes involved in various cellular processes including oxidative stress and inflammation (W.-T. Chen et al. 2020). However, due to technical limitations, the study did not achieve the whole genome scale and single cell resolution at the same time. Here, we adopted a newly developed spatial transcriptomics technology that could achieve both single cell resolution and whole transcriptome coverage, SpaTial Enhanced REsolution Omics-sequencing (Stereo-seq) (A. Chen et al. 2022; Wei et al. 2022), to profile a 6-month-old male 5xFAD mouse. Neurons, as the primary targets of AD, experience both direct and indirect effects from amyloid accumulation over time. Consequently, our research aims to explore transcriptomic changes in neuronal populations in relation to amyloid pathology. We have developed an integrative approach combining image- and non-image-based cell segmentation, non-linear brain section alignment using VoxelMorph, and Unet-based object detection. By analyzing spatially resolved single-cell transcriptomics, we can examine the potential association between transcriptomic alterations in diverse cell clusters and amyloid deposition. Our findings highlight the potential of Stereo-seq technology as a valuable tool for investigating complex neurological disorders.

## Results

### Sample preparation for spatial transcriptomics

The brain of a 6-month-old male 5xFAD mouse was collected and subjected to a tissue preparation protocol. The brain tissue was frozen and embedded in OCT (optimal cutting temperature) medium. After evaluating the quality of the RNA, multiple coronal sections were obtained through cryo-sectioning (**Figure 1A**). The brain sections were then placed on the surface of STOmics chips. Fixation and permeabilization were then applied, allowing for the capture of released mRNA molecules by spatially barcoded probes on the surface of the STOmics chip. Reverse transcription was performed to generate the cDNA library, followed by sequencing (**Figure 1B**). Meanwhile, to determine the correlation between spatial transcriptomic changes and A-beta deposits, we performed A-beta immunostaining on adjacent brain sections (**Figure 1C**). After sequencing, quality control measures were taken to ensure the accuracy of the sequenced data. GRCm38 mouse genome was used as the reference for alignment. In total, we obtained 4.78 billion reads from this sample. After the quality control, the coronal section (hemisphere) contained around 82.3 million effective reads. Based on the barcoded x and y coordinates of captured individual mRNA molecules in the tissue section, we could reconstruct the spatial profile of these mRNA molecules as the spot-by-gene matrix (**Figure 1D**). Then, the image of immunofluorescence (IF) staining for A-beta and the reconstructed spatial transcriptomics would be aligned (**Figure 1E**) for co-analysis.

**Figure 1.**
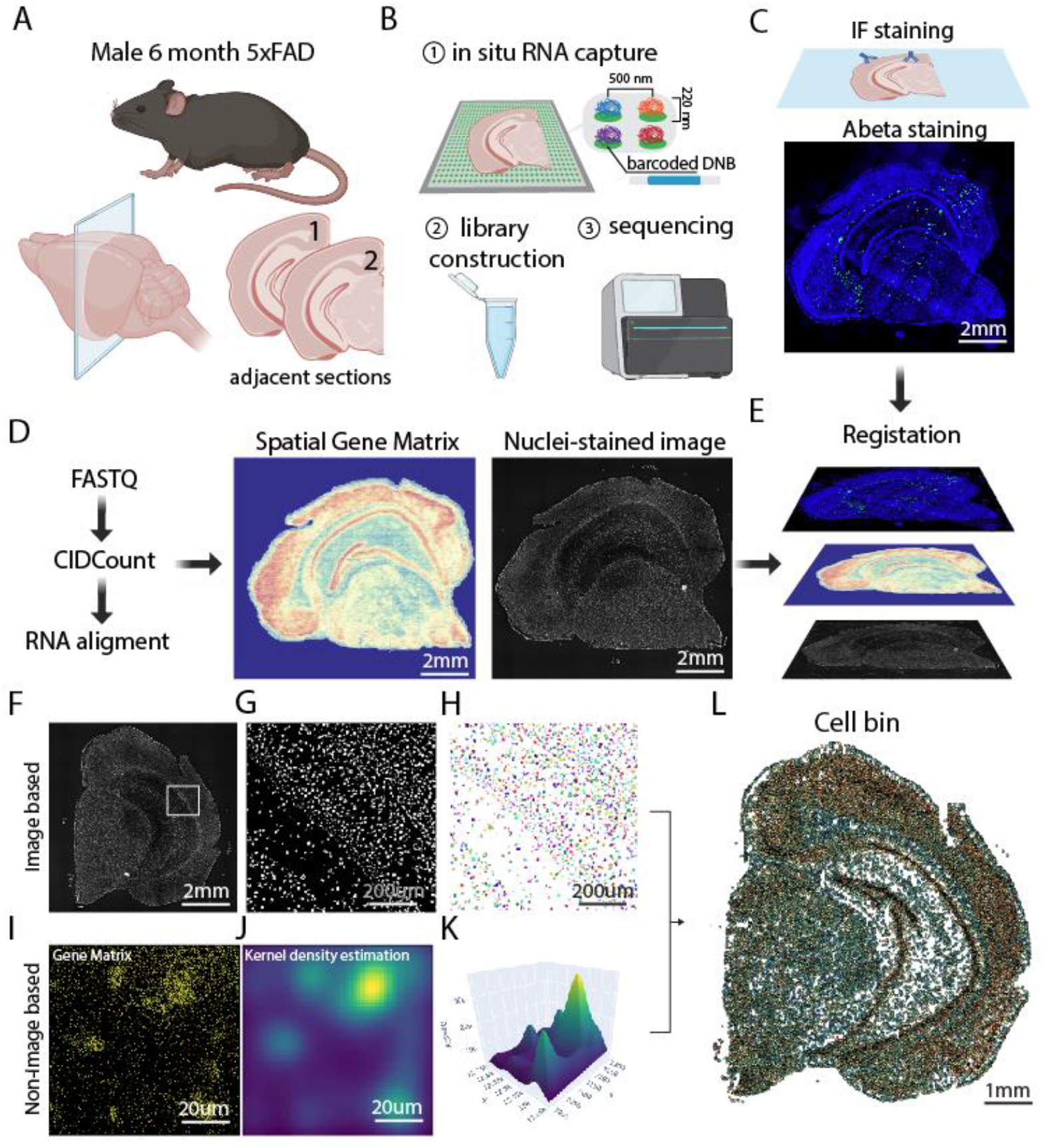
Sample preparation pipeline and cell bin segmentation. **(A)** Two adjacent coronal hemisphere sections obtained from a male 6-month-old 5xFAD mouse. **(B)** One coronal section went through the StereoSeq process to reconstruct the sequencing-based spatial transcriptome. **(C)** The other coronal section went through immunofluorescence staining to visualize the amyloid beta deposition. **(D)** Post-sequencing processing to reconstruct the spatial transcriptomic profile. **(E)** The amyloid beta immunostaining data, spatial transcriptomic data and the ssDNA imaging data were co-registered. (**F-H**) The image-based approach to segment cell bin based on the co-registered ssDNA image. The raw image is shown in (**F**), with (**G**) providing the zoomed-in view. (**H**) shows the instance segmentation of all the nuclei. (**I-K**) The non-image-based approach to estimate the cell location based on the spatial profile of captured RNA density. (**I**) shows the raw spatial distribution of RNA, each dot represents a single RNA copy. (**J**) is the fitted spatial density heatmap using kernel density estimation. (**K**) is the 3D mesh view of the estimated spatial distribution. (**L**) Segmented cell bin through the integration of both image- and non-image-based cell bin segmentation approaches.

### Integrative image- and non-image-based cell bin segmentation

In principle, this gene spatial matrix provides individual transcript information at subcellular resolution, which however requires reliable cell segmentation. Therefore, the generation of a cell bin dataset through single cell segmentation is a crucial step. In this study, we employed an approach that leverages both image- and non-image-based estimation of cell bin. The image-based pipeline consisted of three steps: pixel classification, watershed segmentation, and instance segmentation. The non-image-based pipeline utilized the spatial density estimation and fitting to find the locations of the estimated cells. Then, the image- and non-imaged-based segmentations would be merged to determine the cell bin segmentation (**Figure 1F-K**). In the image-based approach, we utilized the ilastik (Berg et al. 2019) to perform pixel-level classification on the raw single-stranded DNA (ssDNA) image (**Figure 1F-H**). The goal was to distinguish between the cells and the background based on the ssDNA intensity. The output of this step was a binary mask that identified the regions of the image corresponding to the cells (**Figure 1G**). Next, we applied the watershed algorithm to separate the crowded objects in the binary mask. The watershed algorithm is a well-established technique for object separation in images, and it was used in this study to separate cells that were touching or overlapping. The output of this step was an image where each cell was separated from its neighbors and assigned a unique label. This step involved computing the contours of each cell in the label image and plotting them as separate, filled polygons in different colors (**Figure 1H**). We noticed that not all cells indicated by the ssDNA image could be captured, thus underlining the potential problems for solely image-based approaches. The non-image-based approach was based on the property that the RNA molecules tend to aggregate close to the soma (**Figure 1I**). We used the spatial density estimation based on the RNA spatial distribution and found that this could be advantageous in segmenting cells (**Figure 1J, K**). We integrated results from both approaches and only obtained cell bin when two estimation results matched. This cell bin segmentation allowed us to obtain the cell-by-gene matrix for the follow-up analysis.

### Neuronal cell bin clustering

After establishing the cell bin matrix, we took measures of quality control including removing of low-quality cells (with gene counts < 100) and inspection of the spatial distribution of transcript counts (**Figure 2A, B**). In this study, since we focused on the neuronal population, we filtered the dataset using neuronal markers (e.g., Slc17a7, Slc32a1, Sst, Th, etc.). Then, we performed dimensional reduction (based on variable genes) and unbiased clustering using the Louvain algorithm. After the clustering, we visualized the results using a t-SNE (t-Distributed Stochastic Neighbor Embedding) plot (**Figure 2C**). Meanwhile, we obtained a marker gene list for each cluster based on the differential expression analysis (**Figure 2D**). The preservation of the coordinate information of the captured RNA molecules provided us with the opportunity to reconstruct the spatial profile of gene expression. Therefore, we projected the cluster identities to the spatial location (**Figure 2E**). From this spatial plot, we could clearly see clusters localizing in the thalamus (**Figure 2F**), neocortical laminae (**Figure 2G**), and the ventral tegmental area (**Figure 2H**). Cell-type matching was applied to assign reference scRNA-seq cell types to each segmented cell with an associated confidence score based on the identified marker genes as well as their spatial localization. Applying the cell-type matching algorithms produced a cell-by-type matrix as a primary output, consisting of probabilistic assignment of each segmented cell to one of the reference cell types at the subclass level.

**Figure 2.**
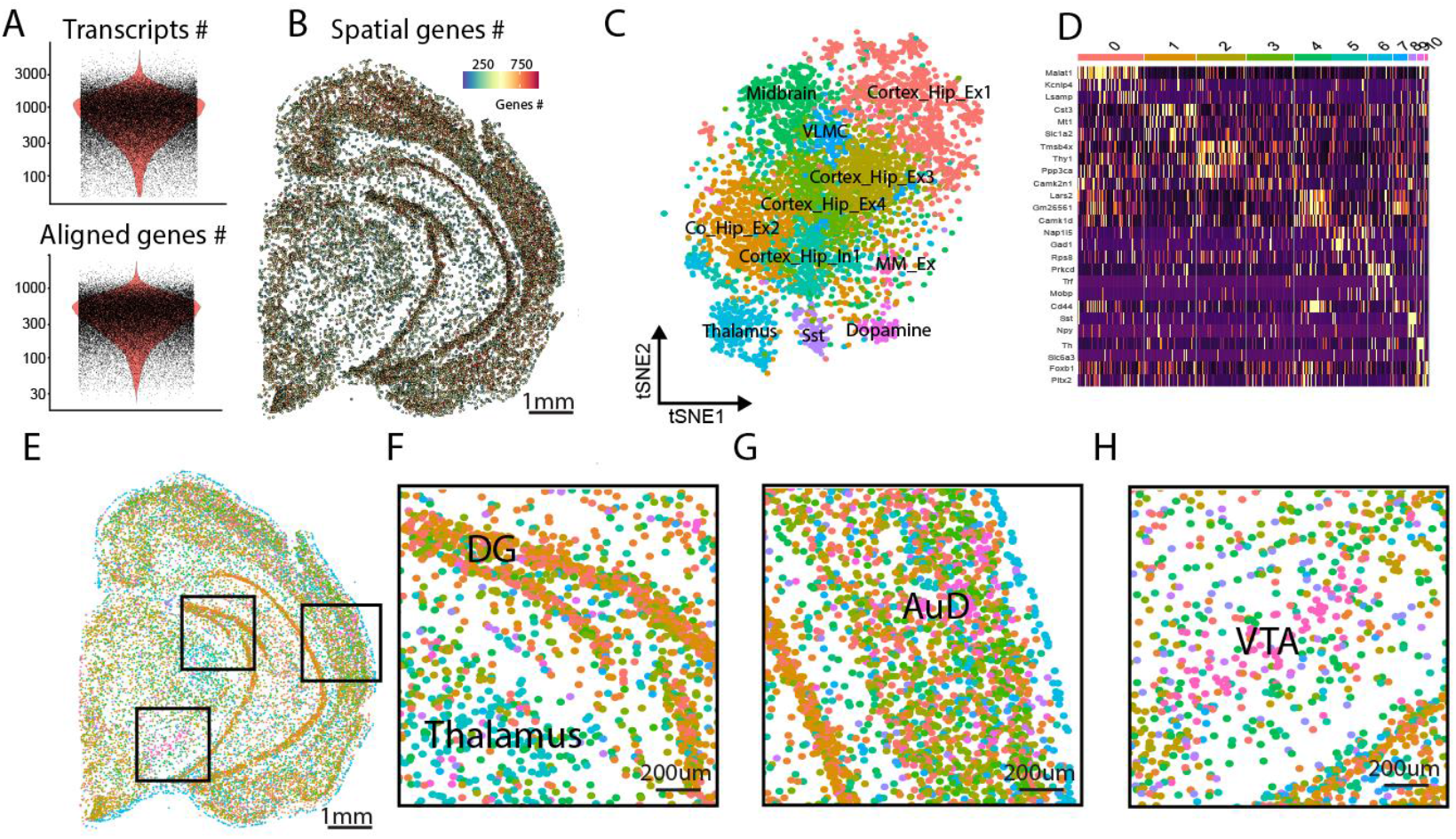
Neuronal cell-type clustering of spatial transcriptomic data. **(A)** Violin plot shows the number of transcripts and aligned genes per cell bin. **(B)** Spatial heatmap of number of genes across the coronal section. **(C)** Dimensional plot of the data using tSNE plot. **(D)** Heatmap shows the marker genes for each cluster (labeled by one color on top). **(E)** Spatial plot of clusters for the whole coronal section. (**F-H**) Zoomed-in view for selected regions in (**E**). DG, dentate gyrus. AUD, auditory cortex. VTA, ventral tegmental area.

### Cross validation of the Stereo-seq data with ISH staining

To evaluate the reliability of the Stereo-seq data, we compared the spatial expression of selected genes reconstructed using the Stereo-seq to that of the *in situ* hybridization (ISH) staining result of the Allen Brain Atlas (**Figure 3**). Prox1 (Prospero Homeobox 1) is a transcription factor with primary expression observed in the dentate gyrus of the hippocampus. Dkk3 (Dickkopf-related protein 3) exhibits expression in the hippocampal CA1-3 regions. Dcn (Decorin) is found in the ventral hippocampal region, while Prkcd (Protein Kinase C Delta) is expressed in the thalamus. Slc6a3 (Solute carrier family 6 member 3), also known as DAT (Dopamine transporter), is predominantly located in the substantia nigra and ventral tegmental area. Lastly, Nrgn (Neurogranin) displays high-level expression in regions relevant to learning and memory, such as the cortex, hippocampus, and amygdala (Lein et al. 2007). The comparison revealed that the spatial expression patterns obtained from the two datasets were highly consistent with each other. This provides further support for the reliability of the Stereo-seq data, demonstrating its ability to accurately capture and reconstruct the spatial gene expression patterns in the tissue.

**Figure 3.**
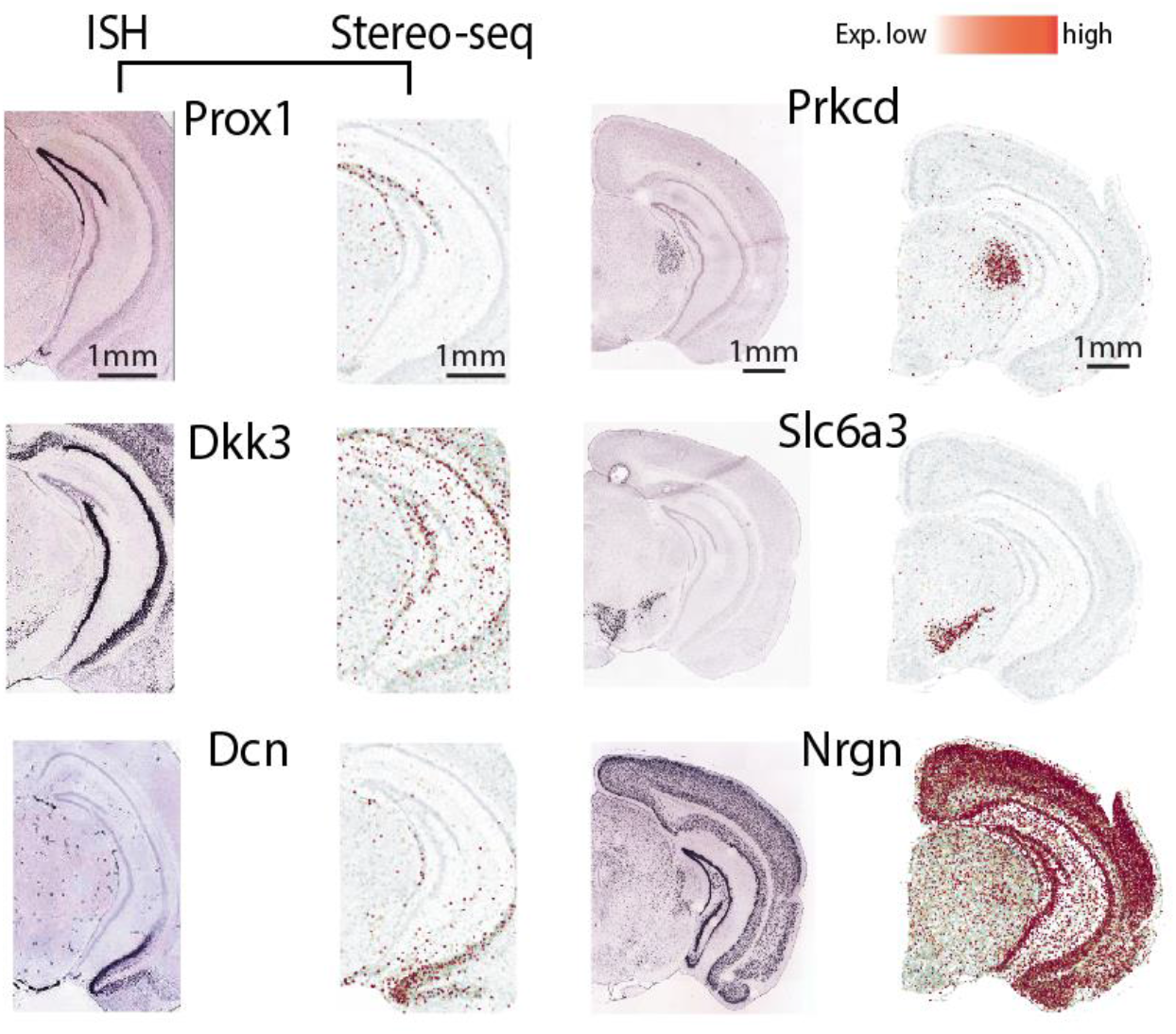
Cross validation with *in situ* hybridization data. Left column shows the ISH image of selected genes, right column shows the spatial distribution of corresponding genes based on the Stereo-seq analysis.

### Amyloid deposition analysis

Using immunohistological staining, we could visualize the accumulation of amyloid plaques in different brain regions (**Figure 4A, B**). We adopted a Unet-based segmentation model to classify pixel and segment the amyloid beta deposit (**Figure 4C, D**). Then we performed non-linear and 3D alignment to register the coronal sections to the Allen Brain Atlas (2017 version, **Figure 4E-G**). From this model output, we could clearly see that amyloid beta deposits exhibit a specific rather than a diffused spatial distribution pattern. Based on the atlas registration, we further quantified the A-beta load in each brain region. Consistent with a previous report (Gail Canter et al. 2019), we observed that A-beta exhibits high-level deposition in the subiculum, hippocampus, entorhinal cortex and cortical amygdala region, etc. (**Figure 4H**). This result further suggests the possibility that the amyloid beta deposition could differentially affect different cellular clusters.

**Figure 4.**
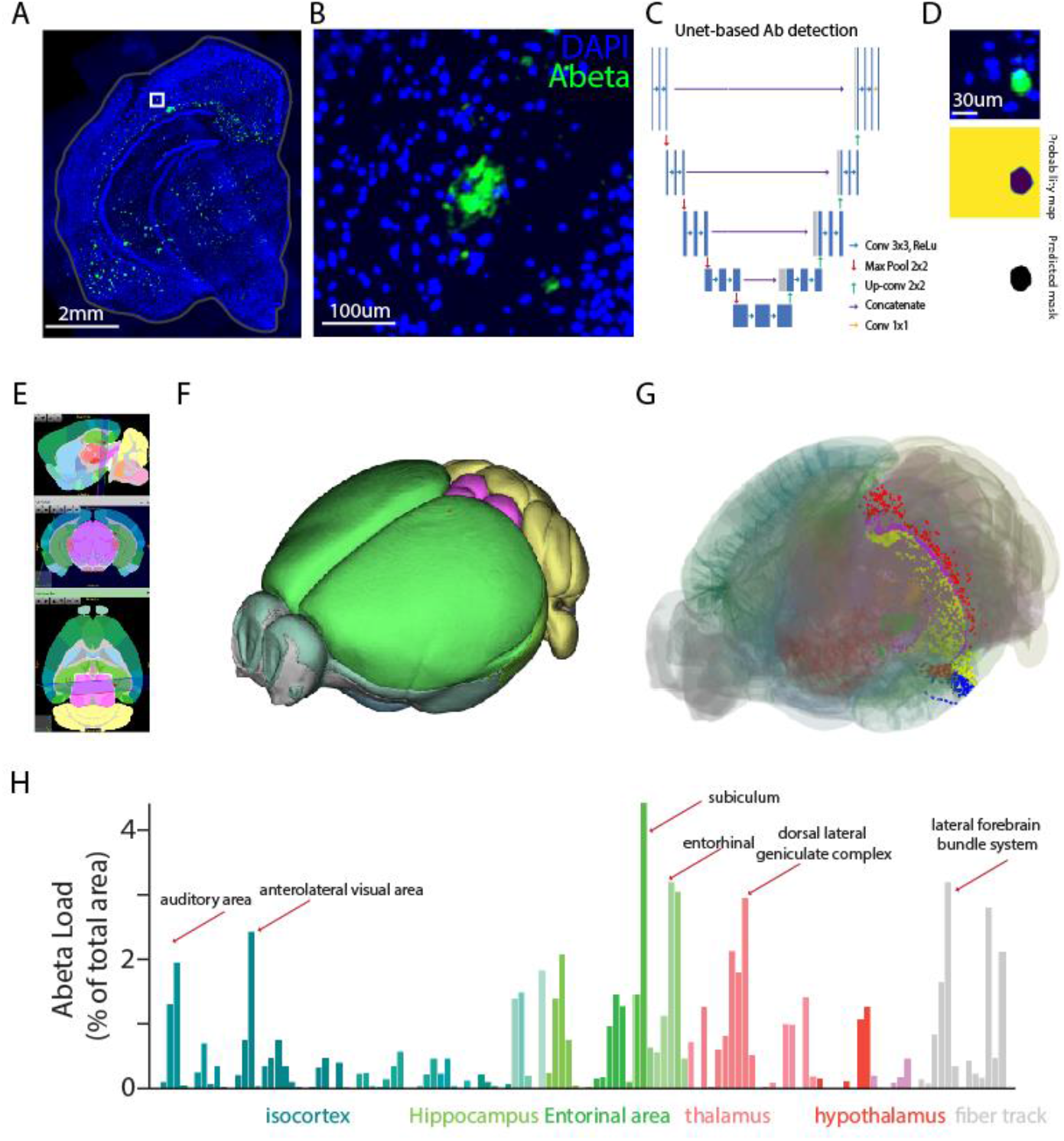
Spatial characterization of amyloid beta deposition. **(A)** Image shows the amyloid beta immunostaining (green) together with DAPI staining (blue). **(B)** Enlarged view of the selected region in (**A**). **(C)** The Unet model architecture for amyloid beta segmentation. **(D)** Example model prediction output of the segmented amyloid beta deposit. **(E)** 3D brain section registration to the Allen Brain Atlas 2017. **(F)** 3D mesh view of mouse brain. **(G)** 3D view of the registered amyloid beta spatial distribution. (**I**) Statistics of amyloid beta load per anatomical area.

### Integrative analysis of spatial transcriptomics and amyloid beta deposition

To correlate the single-cell spatial transcriptomics with the amyloid beta pathohistological map, we co-registered the spatial transcriptomics and the amyloid beta histological data using the Voxel Morph model architecture (**Figure 5A**). Meanwhile, we used the shortest distance to the amyloid beta deposition center as an index to describe the spatial relationship between the cell bin and the amyloid beta plaque (**Figure 5A**). In this way, we could obtain a spatial map of amyloid distance index for each cell bin (**Figure 5B**), from which we could see that some cells were spatially closer to A-beta plaques than others. Next, we applied the amyloid distance index to cell clusters in the dimensional reduction plot and found that some cell clusters are more closely related to amyloid beta plagues in the spatial dimension than others (**Figure 5C-D**), as shown by the ranking of the clusters based on the median amyloid distance index (**Figure 5D**). Using this approach, we would be able to investigate the relationship between cell bin (and cell type) and pathological changes in a spatially resolved manner.

**Figure 5.**
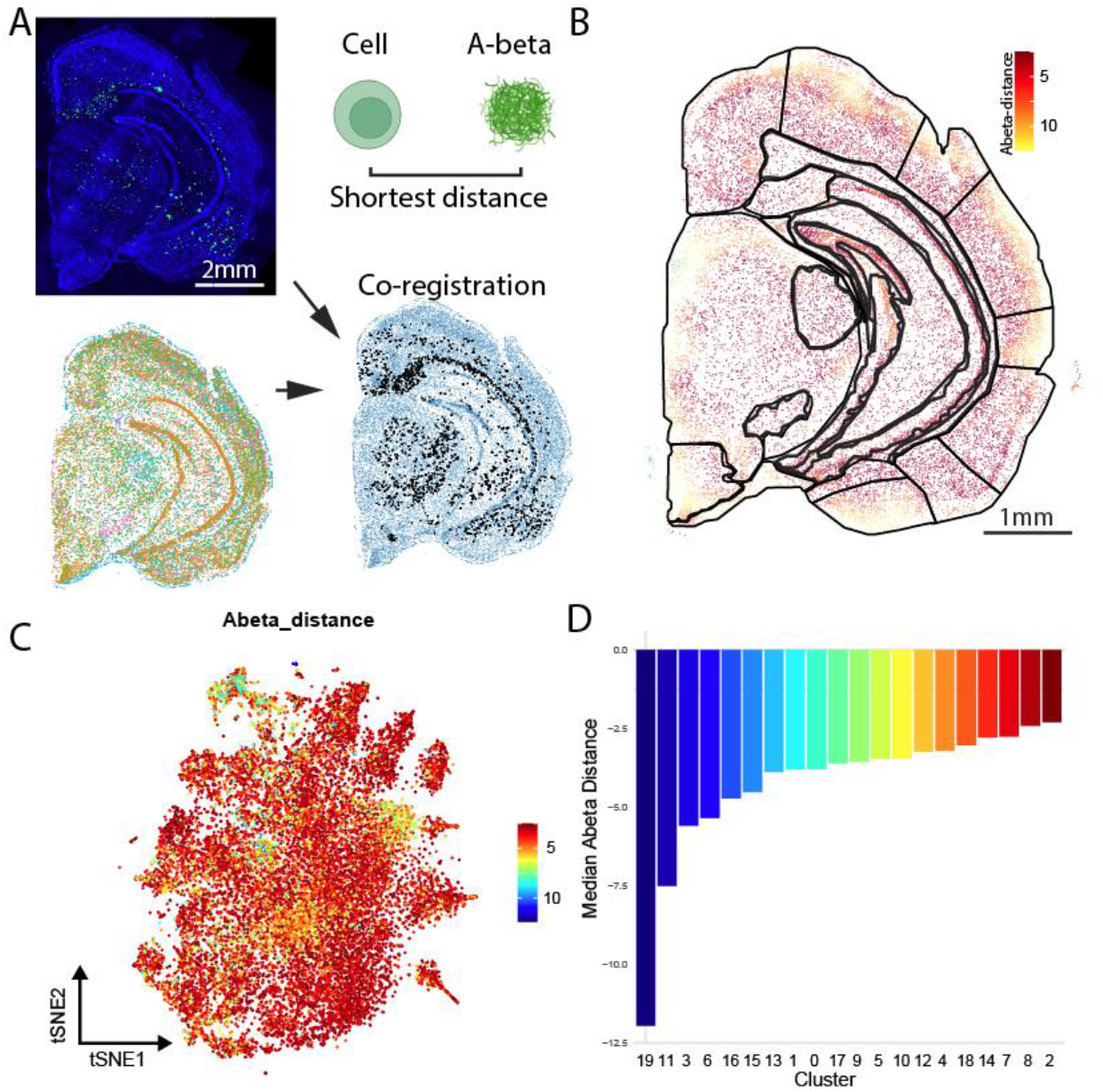
Integrative analysis of neuronal spatial transcriptomics and amyloid beta deposition. **(A)** Nonlinear registration of amyloid beta deposition and spatial transcriptomics data. For each cell bin, the shortest distance towards the center of amyloid beta deposition was calculated. **(B)** Spatial plot of the amyloid distance index for each cell bin. **(C)** Dimensional plot shows the amyloid distance index for each cluster. 1 unit = 50um, **(D)** Ranking of clusters based on median amyloid distance index, with color indicating the relative distance. Blue and red represent the farthest and closest distance respectively.

### Anatomical region-specific analysis

One advantage of using spatial transcriptomics is that it allows us to accurately segment specific anatomical regions for region-specific analysis. We first registered the spatial transcriptomic data to the Allen brain atlas based on visible landmarks using QuickNii (see Methods). Next, we fine-tuned the alignment based on the molecularly defined boundaries for each region (e.g. using thalamus-specific genes to define thalamic boundaries). Based on the integrative anatomy-molecular atlas registration, we could assign the anatomical region tag for each cell bin. To demonstrate the anatomical region-specific analysis, we used the segmentation of the auditory cortical region as an example (**Figure 6A**). After 3D-based atlas registration of the data (**Figure 6A**), we could extract the cell bin data specifically from the anatomically defined region (e.g., auditory cortex), and perform cell-type clustering within the selected region (**Figure 6B-C**). Further, we can visualize their relationship with amyloid beta deposition (**Figure 6D-E**).

**Figure 6.**
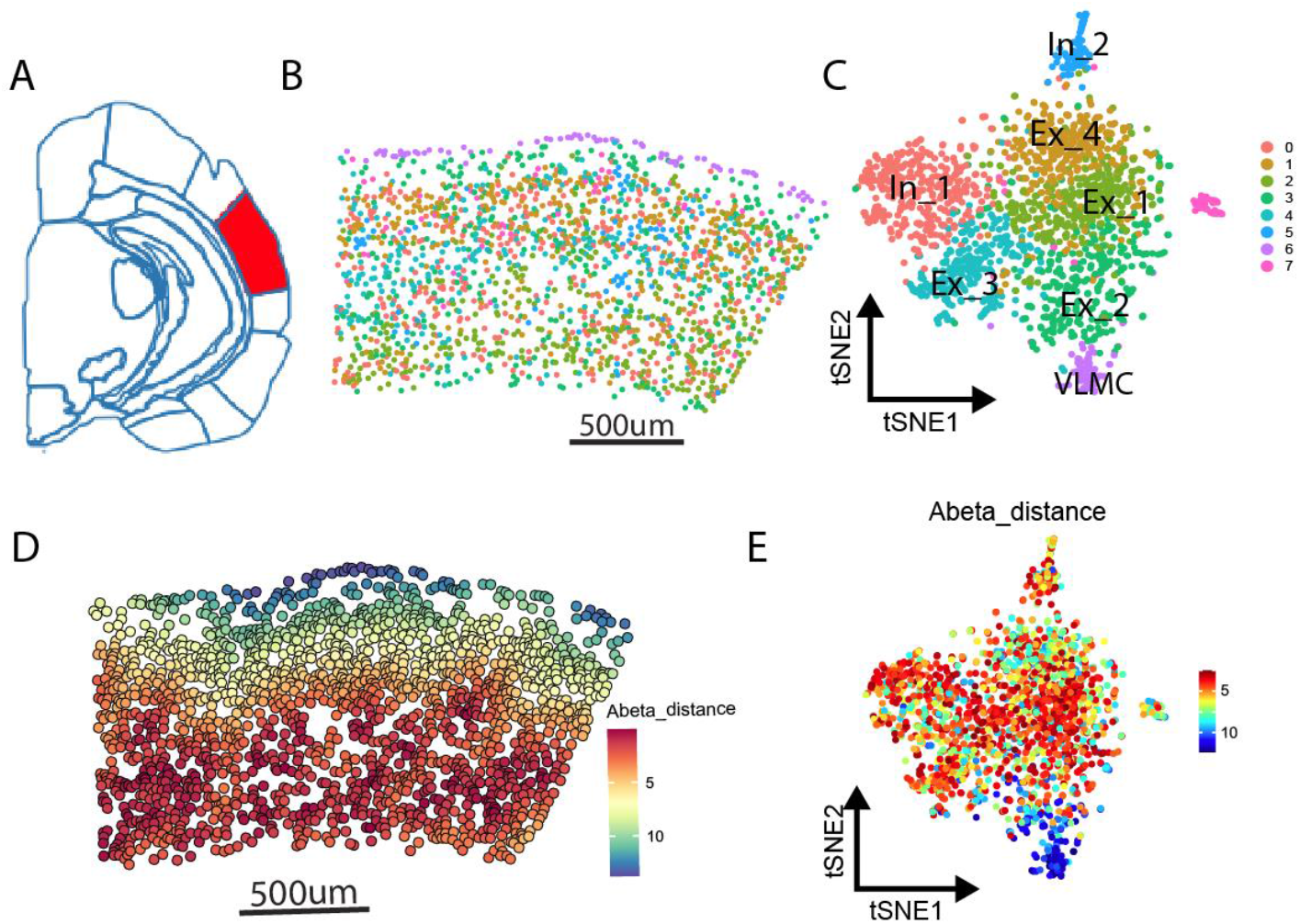
Spatial segmentation analysis of selected anatomical regions. **(A)** The selected anatomical region, covering the auditory cortex (red). **(B)** Spatial plot of cell bin in the selected region, with color representing the identify of clusters. **(C)** Spatial plot of A-beta distance index for each cell bin. **(D)** DimPlot of clusters for the selected region. 1 unit = 50um, **(E)** DimPlot of relative A-beta distance for the selected region.

## Discussion

In this study, we have successfully applied the Stereo-seq technology to analyze transcriptomic profiles in a 6-month-old male 5xFAD mouse model of AD and developed an integrative image- and non-image-based cell segmentation method. By doing so, we could identify vulnerable cell clusters spatially associated with the amyloid-beta deposition. We have validated the reliability of the Stereo-seq data through comparing them to *in situ* hybridization results. In addition, we have performed anatomical region-specific analysis by quantifying the accumulation of A-beta plaques in different brain regions using Unet-based segmentation. By extracting data from selected brain regions with enriched A-beta deposition, we could establish the spatial relationship between cell clusters and amyloid-beta plaques. These results highlight the significance of employing advanced technologies such as Stereo-seq to better understand the molecular mechanisms underlying neurodegenerative disorders such as Alzheimer’s disease.

During this research, we have made significant methodological advancements, particularly with the development of the integrative image- and non-image-based cell segmentation method. The accuracy of the Stereo-seq output based on this cell segmentation method, as assessed by comparing it to *in situ* hybridization results and other ground truth references, highlights the robustness of this technology in capturing and reconstructing spatial gene expression patterns. This further solidifies the potential of Stereo-seq as a valuable tool for investigating complex neurological diseases in general.

As the current focus is on the technique validation, we have only examined a single time point in the AD disease progression. Future studies could benefit from examining multiple time points, which would provide a more comprehensive understanding of the dynamic changes in gene expression patterns and the spatial organization of vulnerable cell populations throughout the disease progression. This information could help to identify key stages in the disease process where interventions might be the most effective. More importantly, the scalability of this technique makes it suitable for application in much larger tissues, including samples from human AD patients.

Another potential direction for future research is the integrative investigation of other neurodegenerative diseases that requires cross-modality approaches, e.g. combined histology and spatial genetic technology. Comparisons between different diseases may then help to identify common and distinct molecular pathways contributing to neurodegeneration, which could inform the development of more targeted and effective therapies.

In conclusion, this study demonstrates the potential of the Stereo-seq technology for investigating complex neurological disorders such as Alzheimer’s disease. Our methodological advancements, in particular the development of an integrative image- and non-image-based cell segmentation approach, provide valuable tools for future spatial transcriptomics studies in neuroscience. The insights gained from the research along this direction can help to improve our understanding of the molecular mechanisms underlying Alzheimer’s disease and other neurodegenerative disorders, paving the way for the development of novel therapeutic strategies.

## Methods

### Animals

We obtained two adjacent coronal sections by cryo-sectioning the brain from a male 5xFAD mouse at 6 months of age. One of the sections was processed for Stereo-seq-based spatial transcriptomic profiling, whereas the other was immunostained for amyloid beta.

### STOmics library preparation

The Omics library preparation was performed as previously described (A. Chen et al. 2022; Wei et al. 2022). Briefly, samples were thawed at -20°C for at least 30 minutes, and section ed coronally using a Leika CM 1950 cryostat (10 μm in thickness). The STOmics chips (1cm x 1cm) were washed with nuclease free water (NF water) (Ambion, Cat. NO. AM9937) twice and dried at 37°C for 1 minute. Then, the brain section was mounted to the surface of the pre-chilled capture chip and incubated at 37°C for 3 minutes. Chips with sections were immersed immediately in pre-cold methanol and incubated for 30 minutes at -20°C before library preparation (see below).

### Single cell nuclei staining

After fixation or immunofluorescence (IF) staining on chips, sections were stained with Qubit ssDNA reagent (Thermo Fisher, Q10212) diluted in 5 × SSC (Thermo Fisher, AM9763) for 5 minutes. Chips were further washed with 0.1x Wash Buffer (0.1 × SSC supplemented with 5% RNAase inhibitor [BGI-Research, 1000028496]) and dried for imaging. The Motic Custom PA53 FS6 microscopy was used for nuclei staining images at the channel of FITC with a 10x objective len.

### Tissue permeabilization

After imaging, tissue section mounted on the chip was permeabilized with PR Enzyme (BGI-Research, 1000028496) in 0.01M HCl buffer at 37°C for 12 to 15 minutes, and washed with 0.1x Wash Buffer.

### In situ reverse transcription (RT)

After permeabilization, the released mRNA molecules that were captured by DNA nanoball (DNB) were reverse transcribed immediately at 42°C for 3 hours using RT Mix (BGI -Research, 1000028496), including RT reagent, RT Additive, RNAase Inhibitor, RT Oligo and Reverse Transcriptase Enzyme.

### Tissue removal

The same chips with brain sections were then washed with 0.1x Wash Buffer and digested with Tissue Removal buffer (1BGI-Research, 1000028496) at 55°C for 10 minutes.

### cDNA purification and amplification

The cDNA Release Mix (BGI-Research, 1000028496), including cDNA Release Enzyme and cDNA release Buffer was added to the chips and incubated at 55°C overnight. The resulting cDNAs were purified with VATHS DNA Clean beads (0.8x, Vazyme, N411-03), and amplified with cDNA Amplification Mix and cDNA Primer (5-CTGCTGACGTACTGAGAGGC-3) (BGI-Research, 1000028496). PCR reactions were set up as follows: incubation at 95°C for 5 minutes, 15 cycles at 98°C for 20 seconds, 58°C for 20 seconds, 72°C for 3 minutes and a final incubation at 72°C for 5 minutes, hold infinite R 12°C.

### STOmics library construction and sequencing

The STOmics library construction and sequencing procedures were conducted as previously described by Chen et al. (2022) and Wei et al. (2022). The resulting PCR products were further purified with VATHS DNA Clean beads (0.6x, Vazyme, N411-03) and quantified by Qubit dsDNA Assay Kit (Thermo, Q32854). A total of 20 ng of DNA was then fragmented with in-house Tn5 transposase at 55°C for 10 minutes, after which the reactions were stopped by the addition of 0.02% SDS and gently mixing at 37°C for 5 minutes. Fragmented products were amplified as follows: 25 ml of fragmentation product, 1 3 KAPA HiFi Hotstart Ready Mix and 0.3 mM Stereo-seq-Library-F primer, 0.3 mM Stereo-seq-Library-R primer in a total volume of 100 ml with the addition of nuclease-free H_2_O. The reaction was then run as: 1 cycle of 95°C 5 minutes, 13 cycles of 98°C 20 seconds, 58°C 20 seconds and 72°C 30 seconds, and 1 cycle of 72°C 5 minutes. PCR products were purified using the AMPure XP Beads (0.63 and 0.153), used for DNB generation and finally sequenced on MGI DNBSEQ-Tx sequencer.

### Adjacent sections with IF staining

The adjacent section was fixed in pre-chilled methanol for 5 minutes at -20°C, and further subjected to 1xDPBS (Gibco, 14190-144) to remove the excessive OCT. After creating hydrophobic barrier with ImmEdge Pen (Vector, H-4000), the section was blocked with 5% donkey serum (Sigma, D9963) blocking buffer at room temperature for 30 minutes, and then incubated with FITC conjugated mouse anti-beta Amyloid (MOAB-2) (Novus, NBP2-13075F) and Hoechst dye (H3954) for 15 mins at room temperature. Afterwards, the section was washed with 1xDPBS for twice, mounted with Antifade Mounting Medium (Vector, H-1000) and imaged with a Nikon confocal microscope.

### Immunostaining of beta amyloid

To determine the correlation between transcriptional changes and A-beta deposits, we performed A-beta immunostaining on the adjacent brain section. The section was first blocked with Blocking Buffer (20xSSC, donkey serum, 10% Triton-x-100, RI, NF water) for 15 minutes on ice, and then incubated with FITC-conjugated mouse anti-beta Amyloid (MOAB-2) (Novus, NBP2-13075F) with the same Blocking Buffer for another 15 minutes on ice. The section was then washed twice with Wash Buffer (20xSSC, 10% Triton-x-100, 5% RI, NF water) and once with 0.1xSSC. Nuclei were stained with DAPI (ThermoFisher, D1306). Afterwards, we aligned the cell bin section with the amyloid-beta/DAPI staining section.

### STOmics raw data processing

The MGI DNBSEQ-Tx sequencer generated the Fastq files. Read 1 consisted of coordinate identity (CID) (1-25bp) and molecular identity (MID) (26-35 bp) sequences, while read 2 contained the sequence of cDNA. The STOmics Analysis Workflow (SAW) was used to perform quality control of sequencing data, genome alignment and quantification of gene expression. Firstly, CID sequences in Read 1 were compared with barcode sequences on STOmics chip, and read pairs containing valid CIDs are extracted. For read pairs containing valid CIDs, the CID sequence was converted into the spatial position information on the slice and written into the read ID of Read 2. Then valid reads in Read 2 were filtered out as final Clean Reads. Clean reads are mapped to the GRCm38 reference genome for alignment, and the number of reads mapped to exon regions, intron regions and intergenic regions was counted. Reads overlapped more than 50% with the exon region were counted as exon transcripts. Reads overlapped less than 50% with the exon region yet possessing overlapped sequences with the adjacent intron sequence were annotated as intron transcripts, otherwise as intergenic transcripts. Finally, the exonic reads were used to generate a CID-containing expression profile matrix. Then, locating Unique Mapping Reads to genes, removing duplicate MIDs, and calculating the expression levels of all genes according to MID correction were performed. The Stereo-seq method is based on DNA nanoball (DNB)-patterned arrays and in situ RNA capture to reconstruct the spatial transcriptomic map. The diameter of the DNA nanoball is 220nm. The center-to-center distance of adjacent DNA nanoballs is 500μm.

### Spatial gene matrix

The spatial transcriptomic data obtained using stereo-Seq is a collection of detected molecules, each corresponding to a specific gene, with predefined and barcoded coordinates of the molecule within the field of view. The gene matrix table contained information regarding each nanoball with gene name, x and y coordinate, as well as the count number.

### Cell bin segmentation

Cell pixel classification was performed using ilastik (Berg et al. 2019) to generate the segmented image. To detect the cell boundary, distance transform watershed was used as instance segmentation. Size opening was performed to reduce background noise, and smaller objects that would not fit into cellular shape were removed.

### Brain section registration

The registration was divided into two steps. First, we applied the linear affine alignment in the 3D space using QuickNii ABAMouse 2017 (https://www.nitrc.org/projects/quicknii). Next, the non-linear deformation 2D fine tuning with intervention was performed by experienced experimenters using the VisuAlign (https://www.nitrc.org/projects/visualign/). The registered colormap could be converted to the polygon shapefile using Adobe Illustrator (https://www.adobe.com) and Qupath (https://qupath.github.io/). The shapefile will be used for the cell bin or 50 bin registration using customized python code (in Python2.8, primarily using the geopandas library, Written by Guang-Wei Zhang, which will be available on Github: https://github.com/GuangWei-Zhang).

### Postprocessing

Seurat V4, stereopy (https://github.com/BGIResearch/stereopy), and spateo (Qiu et al. 2022) were used in this study were used in this study. The h5seurat files for each brain section will be loaded using a customized data loader to match the Seurat format. For the stereopy analysis, the .gef cell bin files were adopted. Detailed scripts that could replicate all results in this study will be made available on Github (https://github.com/GuangWei-Zhang).

### Quantification of A-beta deposition

The amyloid-beta and ssDNA images were registered using nonlinear transformation based on VoxelMorph framework. The customized Unet model was used for amyloid-beta segmentation. The registration and quantification were achieved through the QUINT processing pipeline (Groeneboom et al. 2020).

